# Clinically derived micro- and nanoplastics uptake drives spatiotemporally confined metabolic stress revealed by bond-selective imaging

**DOI:** 10.64898/2026.07.02.735952

**Authors:** Jingyuan Li, Nian Liu, Delong Zhang, Hyeon Jeong Lee

**Author notes:** Corresponding author: Hyeon Jeong Lee. These authors contributed equally: Jingyuan Li, Nian Liu.

## Abstract

Although microplastics and nanoplastics (MP/NP) are pervasive environmental contaminants, our understanding of cellular toxicity remains incomplete, as adverse effects are often attributed to long-term intracellular accumulation, while the spatiotemporal onset of cellular damage remains poorly defined. Here, we employ chemical-bond-selective stimulated Raman scattering (SRS) microscopy and cell models that decouple continuous exposure from intracellular retention to directly visualize clinically derived MP/NP-cell interactions. Cellular stress occurs primarily during MP/NP exposure, accompanied by alterations in lipid droplet (LD) composition. In contrast, following extracellular removal, intracellularly retained MP/NP become largely inert, with recovery of lipid metabolism and cellular functions. Lipidomics identifies arachidonic acid (AA) as a key dysregulated metabolite, and SRS imaging further reveals transient, spatially confined AA enrichment in MP/NP-proximal LDs during uptake. Importantly, phospholipid coating of MP/NP attenuates LD alterations and cytotoxicity while preserving particle internalization, establishing uptake-driven metabolic stress, rather than long-term intracellular retention, as primary source of MP/NP-induced damage.

## Introduction

Increasing evidence of micro- and nanoplastic (MP/NP) accumulation in human tissues and biofluids [1–5] has brought growing attention to their potential health risks [6–8], yet the biological mechanisms by which MP/NP interact with cells and cause toxicity remain unclear [9, 10]. In vivo studies have reported diverse biological consequences of MP/NP exposure across multiple organs, including the liver [11–13] and cardiovascular system [7, 14], ranging from tissue-level dysfunction [3] to systemic pathophysiological effects [15]. At the cellular level, MP/NP have been associated with oxidative stress [16, 17] and metabolic perturbations [18, 19], although the reported effects vary substantially across studies [11, 20]. Meanwhile, accumulating evidence suggests that MP/NP exposure perturbs fundamental cellular metabolic processes, with potential broad and long-lasting physiological consequences. For example, exposure to polystyrene microplastics has been shown to inhibit monosaccharide metabolism and induce abnormal fatty acid accumulation in the liver [21], and biodegradable polylactic acid microplastics can exacerbate lipid metabolic disorders and even induce cardiac dysfunction through activation of the PPARγ pathway [22]. Together, the substantial variability across studies suggests that MP/NP-induced effects cannot be fully explained by bulk or endpoint analyses, highlighting the importance of mechanistic investigations at the subcellular level.

Given that MP/NP are chemically stable and largely resistant to degradation [23] a central but underexplored question is how and at which stage cellular damage arises, whether from continuous extracellular exposure to MP/NP, from the process of cellular uptake, or from long-term intracellular particle retention. Most existing studies assess toxicity using endpoint assays and do not distinguish among these phases of MP/NP–cell interaction. Moreover, mammalian toxicity studies commonly employ manufactured microspheres with uniform size, shape, and surface properties, often on the order of several micrometers in diameter (e.g., 5 μm) [20]. While such models facilitate controlled exposure studies, they do not capture the physical and morphological complexity of environmentally derived MP/NP.

At the same time, naturally derived MP/NP lack intrinsic optical contrast and are therefore inherently difficult to visualize and track within cells. Although fluorescence labeling is commonly used to monitor MP/NP uptake, such labeling can perturb native cellular physiology through unintended photochemical reactions or phototoxicity [24–26]. Together, these challenges highlight the critical need for label-free imaging approaches capable of directly visualizing naturally derived MP/NP uptake dynamics and intracellular distribution in living biological systems.

For this purpose, vibrational spectroscopic approaches, including Raman and IR, have demonstrated the feasibility of MP/NP detection in biological samples. Leveraging intrinsic chemical bond selectivity, spontaneous Raman spectroscopy has been used to identify MPs in human urine and kidney tissues without labels [5], providing direct evidence of MP presence in human systems. To further improve imaging speed and throughput, stimulated Raman scattering (SRS) microscopy has been developed, enabling submicron-resolution imaging of specific molecular vibrations [27–29]. More recently, SRS microscopy has been successfully applied to investigate MP/NP in bottled water [30] and in vivo monitoring of MP/NP [31], demonstrating its capability for chemically specific imaging of MPs in solution and complex biological environments.

Here, we employ chemical-bond-selective SRS microscopy to directly visualize interactions between clinically derived MP/NP and intracellular organelles to investigate associated metabolic alterations. Hyperspectral SRS imaging of cells continuously exposed to MP/NP reveals pronounced alterations in lipid droplet (LD) composition. In contrast, once extracellular MP/NP are removed, intracellularly retained MP/NP become largely metabolically inert, indicated by recovery of lipid metabolism and cellular functions. Untargeted lipidomics further demonstrates extensive lipid metabolic remodeling under continuous exposure, identifying arachidonic acid (AA) as a key dysregulated lipid species. Importantly, through SRS imaging of individual LDs and MP/NP, we uncover localized enrichment of AA in LDs proximal to MP/NP during uptake, forming a transient and spatially confined AA elevation that dissipates following extracellular MP/NP removal. The uptake-associated mechanism is further supported by the observation that phospholipid coating of MP/NP effectively attenuated LD composition changes and cytotoxic responses while preserving MP/NP internalization, implicating surface-mediated interactions during uptake as an important driver of MP/NP-induced cellular damage. Overall, these findings indicate that freshly internalized MP/NP drive metabolic dysregulation and cellular stress, whereas long-term intracellular particle retention plays a minimal role.

## Results

### Tracking MP/NP uptake and distribution in live cells by SRS imaging

To simulate the complex physical morphology of MP/NP in natural environments, MP/NP used in this study were clinically derived, originated from medical-grade bone cement used in implants (see Methods), and these clinically derived MP/NP were subjected to morphological and chemical characterization using scanning electron microscopy (SEM) and SRS microscopy (Fig. S1). SEM analysis revealed that the isolated MP/NP exhibited predominantly irregular, near-spherical morphologies with diameters ranging from 200 to 3500 nm (Fig. S2A, B), and the surface charge of the MP/NP is -38 mV (Fig. S2C). To characterize the chemical signatures and validate the capability of SRS microscopy for MP/NP detection, particles dispersed in water were imaged at the C−H stretching frequency (2950 cm^-1^), demonstrating sufficient spatial resolution for single-particle visualization (Fig. S2D). Chemical composition was further analyzed through hyperspectral SRS imaging across 2800 to 3100 cm^-1^. The spectrum showed characteristic peaks at 2950 cm^-1^ (aliphatic −CH2− stretching) and 2850 cm^-1^ (symmetric C−H stretching), consistent with the known vibrational spectrum of polymethyl methacrylate (PMMA) [32] (Fig. S2E). To validate the ability of SRS imaging to detect internalized particles, MP/NP were added to the culture medium and imaged at 2950 cm^-1^. Aggregates were observed within cells (Fig. S2F), and size analysis on individually dispersed intracellular particles confirmed the presence of both micrometer- and nanometer-sized plastics (Fig. S2G, H).

To investigate the temporal and spatial dynamics of MP/NP interactions with cells, we established two complementary cell models (Fig. 1A). In the continuous-exposure model, cells were maintained in MP/NP-containing medium and imaged by SRS at multiple time points up to 72 hours (T0, T2, T24, and T72), enabling characterization of cellular responses while MP/NP remained constantly present in the extracellular environment. In parallel, a washout model was established where MP/NP-containing medium was replaced with MP/NP-free medium after initial exposure. SRS imaging was then performed at corresponding time points up to 72 hours (T0; T24: 24 h exposure; T24-W24: 24 h exposure followed by 24 h withdrawal; T24-W72: 24 h exposure followed by 72 h withdrawal), thereby isolating the effects driven solely by intracellularly retained MP/NP. Phasor analysis in the C–H region (2800 to 3100 cm⁻¹) decomposed hyperspectral SRS datasets into four components: lipid, nucleic acid, cytoplasm, and MP/NP (Fig. S3). The MP/NP-specific component (ROI 5) appeared only in MP/NP-exposed cells, demonstrating the high chemical specificity of hyperspectral SRS combined with phasor analysis for intracellular MP/NP tracking.

**Fig. 1.**
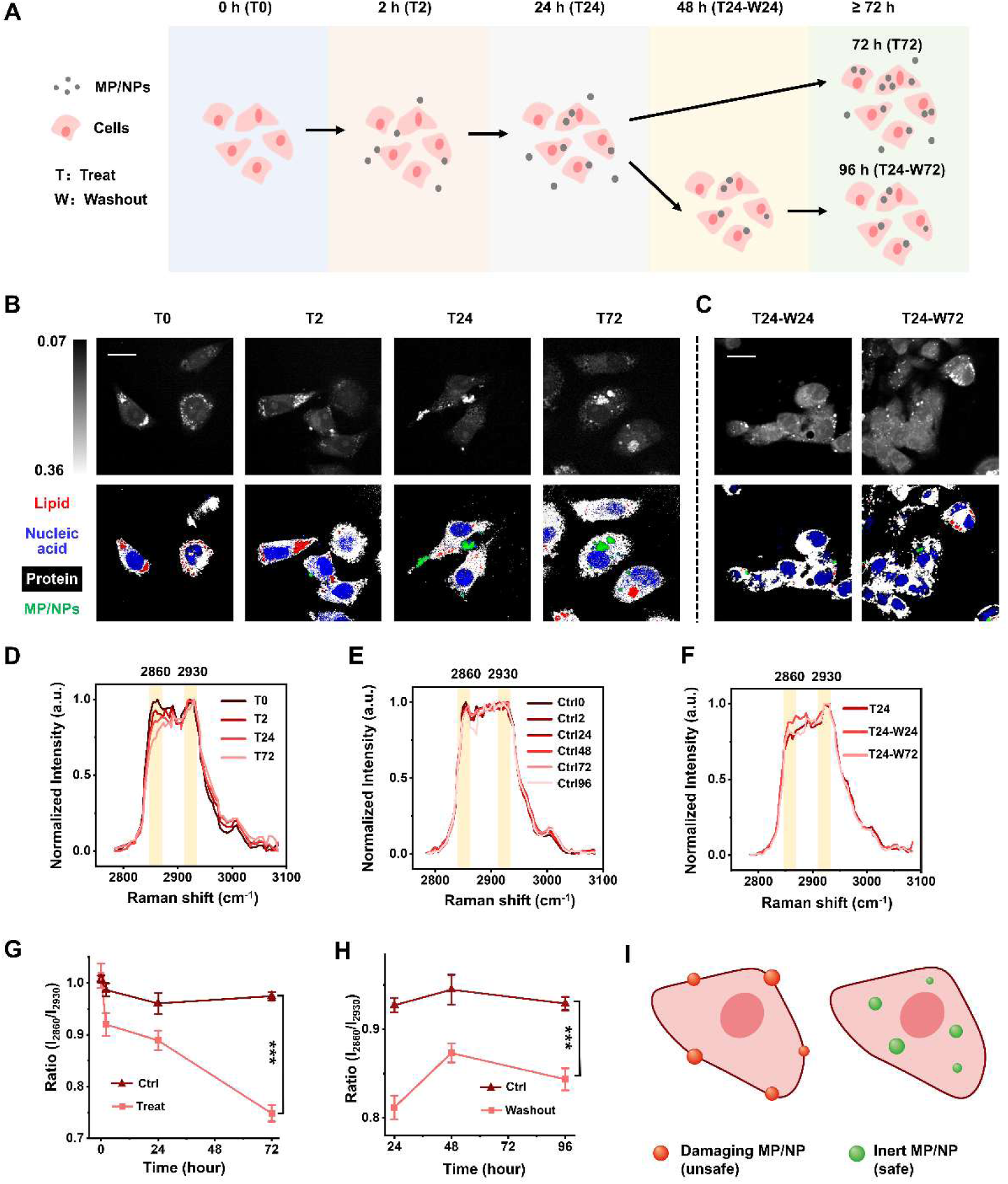
Continuously treat MP/NP induced remodeling of LD composition in HepG2 cells while it can be recovered with MP/NP washout in medium. **(A)** Schematic diagram of the experimental timeline for treating cells with MP/NP. (**B** and **C**) Representative SRS images at 2913 cm^-1^ and corresponding merged images from phasor analysis. Red, blue, white, and green represent lipid, nucleic acid, protein and MP/NP respectively. Scale bar, 10 μm. (**D**) SRS spectra at 2800-3100 cm^-1^ of LDs extracted from MP/NP-treated cells in T0, T2, T24 and T72 groups. (**E**) SRS spectra at 2800-3100 cm^-1^ of LDs extracted from control cells in Ctrl0, Ctrl2, Ctrl24, Ctrl48, Ctrl72 and Ctrl96 groups. (**F**) SRS spectra at 2800-3100 cm^-1^ of LDs extracted from MP/NP-treated cells in T24, T24-W24 and T24-W72 groups. (**G**) The ratio of I_2860_/I_2930_ from the SRS spectra comparing MP/NP-treated cells (red line) and control cells (brown line). (**H**) The ratio of I_2860_/I_2930_ from the SRS spectra comparing MP/NP-washout cells (red line) and control cells (brown line). (**I**) Diagram of the cell continuously treat with MP/NP or the cell washout with MP/NP. The characteristic peaks are labeled and highlighted. Values are mean ± SD (n=3). *** indicate P < 0.001.

We then analyzed the dynamic internalization process of MP/NP across time points (Fig. 1B). At 2 h of continuous exposure (T2, 10 μg/mL), MP/NP were primarily localized near the plasma membrane. Notably, higher exposure concentrations resulted in increased internalization within the same time frame (Fig. S4), demonstrating the importance of applying physiologically relevant concentrations (∼12 μg/mL in human blood [33]) for evaluating the uptake kinetics. By 24 h (T24), extensive internalization had occurred, with particles distributed throughout the cytoplasm. By 72 h (T72), MP/NP aggregation became apparent, forming clustered subcellular structures. In contrast, the spatial distribution of MP/NP in the washout group (T24-W24 and T24-W72) showed no significant differences compared to T24 (Fig. 1C). To quantitatively assess these dynamics, the number of intracellular MP/NP per cell was quantified. During continuous exposure, the number of intracellular MP/NP increased steadily. However, following the washout, the number remained stable, confirming the long-term retention of MP/NP after internalization (Fig. S5). To further elucidate the internalization mechanism, we inhibited major uptake pathways. When pinocytosis was inhibited by EIPA, MP/NP uptake was significantly reduced, whereas the endocytosis inhibitor Chlorpromazine induced no significant change, suggesting that pinocytosis is the primary internalization mechanism (Fig. S6). These results collectively demonstrate the dynamic internalization and intracellular distribution of MP/NP and provide direct visualization of their temporal and spatial progression in living cells.

### Continuous MP/NP exposure alters LD composition and induces reversible cytotoxicity in cells

Using this complementary cell model that defines MP/NP internalization dynamics, we next assessed the metabolic and functional consequences of MP/NP exposure. Because LDs play a central role in lipid homeostasis, we first examined whether MP/NP exposure perturbs LD composition. Hyperspectral SRS analysis revealed substantial time-dependent alterations in LD spectral signatures under continuous MP/NP exposure. MP/NP-exposed cells exhibited progressively reduced 2860/2930 cm⁻¹ peak intensity ratios (Fig. 1D; offset spectra shown in Fig. S7), whereas MP/NP-free control cells maintained stable spectral profiles across all time points (Fig. 1E). Notably, LD spectral profiles showed substantial recovery following extracellular MP/NP removal (Fig. 1F). Quantitative analysis further validated a gradual reduction in the 2860/2930 ratio during continuous MP/NP exposure, reaching 1.35-fold reduction at 72 h compared with baseline (Fig. 1G). Upon removal of MP/NP from the medium, LD spectral profiles began to recover, with the 2860/2930 ratio increasing 1.18-fold relative to the 24 h exposure group, indicating a partial restoration of LD spectral profile toward MP/NP-free controls (Fig. 1H). Cells cultured in normal medium showed stable 2860/2930 ratios (Fig. 1G, H, 1.00 ± 0.04), suggesting the metabolic perturbation induced specifically by environmental MP/NP exposure. These findings demonstrate that LD compositional alterations occur only during sustained exposure to extracellular MP/NP and are reversible when exposure ceases, even though intracellular MP/NP remain.

We then evaluated whether this metabolic perturbation is accompanied by cellular stress and functional impairment. Live/Dead staining and CCK-8 assays showed a significant decline in viability under continuous MP/NP exposure, whereas extracellular MP/NP removal led to a notable recovery (Fig. 2A, B). Intracellular reactive oxygen species (ROS) levels exhibited a sustained increase during continuous MP/NP exposure, but the increase plateaued and modestly declined following extracellular MP/NP removal (Fig. 2C, D). Immunofluorescence staining of CYP3A4, a key metabolic enzyme in hepatocytes, revealed a time-dependent upregulation during continuous MP/NP exposure, with the highest expression at 72 h, suggesting a cellular defense response to the foreign particles [34, 35]. CYP3A4 levels decreased after extracellular MP/NP removal, most prominently at 72 h post removal (Fig. 2E, F). Furthermore, the secretory function of hepatocytes was significantly impaired under continuous MP/NP exposure, as evidenced by progressively decreased urea and albumin in the culture medium (Fig. 2G). This impairment began to recover once extracellular MP/NP were removed (Fig. 2G).

**Fig. 2.**
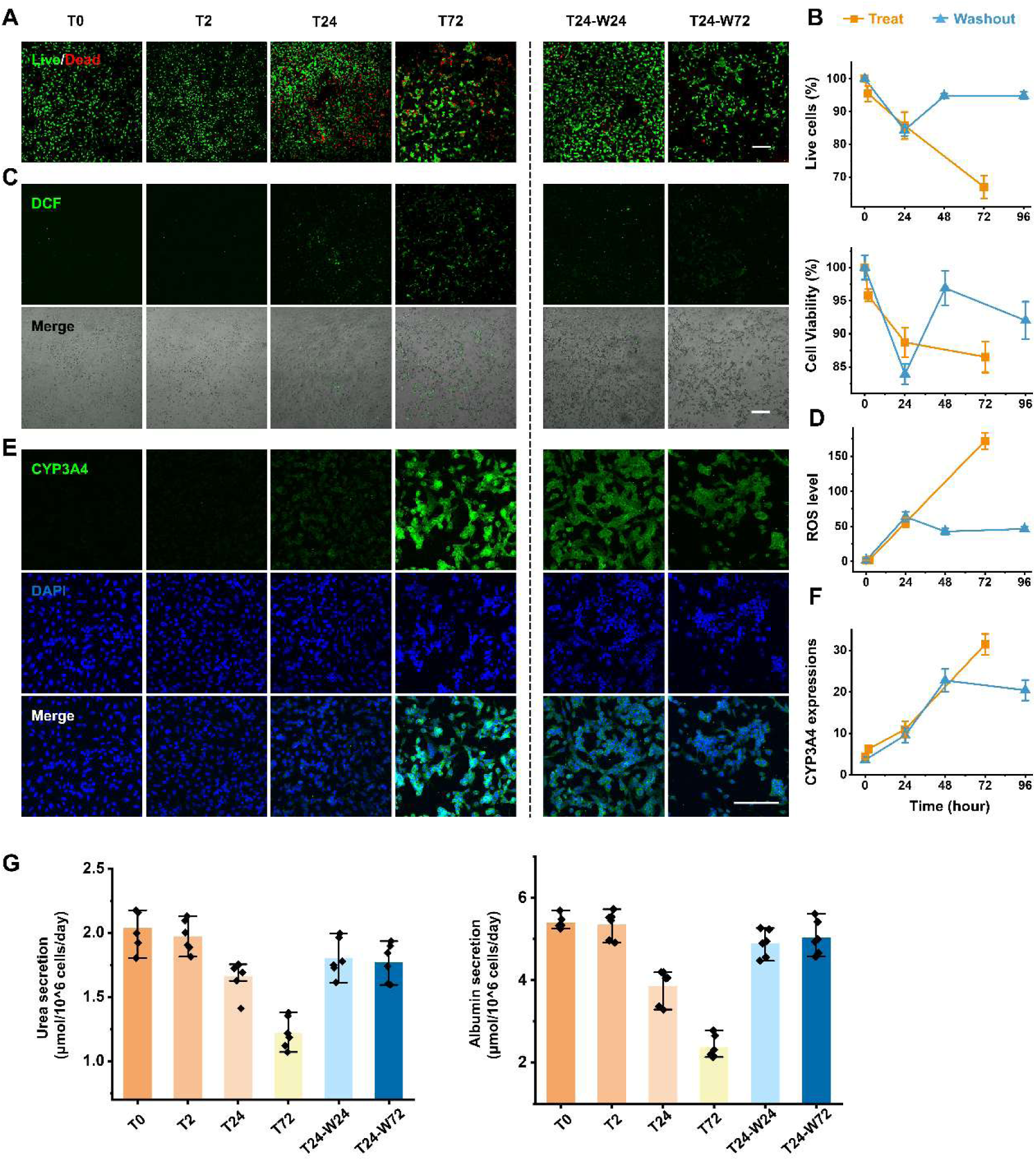
Functional assays of continuously MP/NP-exposed and MP/NP-washout HepG2 cells. (**A**) Representative Live/Dead fluorescence images with green representing live cells and red representing dead cells. (**B**) Live cells percentage from the Live/Dead staining assay and cell viability evaluated by CCK-8 assay in MP/NP-treat and MP/NP-washout groups. (**C**) Representative fluorescence and bright-field merged images, labeled by DCFH-DA. (**D**) ROS levels in MP/NP-treat and MP/NP-washout groups cells. (**E**) Representative immunofluorescence images of CYP3A4 protein. (**F**) Quantification of CYP3A4 expressions. (**G**) Urea and albumin secretion were examined by ELISA kit. Values are mean ± SD (n=3). Scale bar, 400 μm.

Taken together, these results suggest that continuous extracellular MP/NP exposure alters LD composition, induces cytotoxicity, elevates oxidative stress, and impairs hepatocyte function, whereas extracellular MP/NP removal enables substantial metabolic and functional recovery. Importantly, the absence of further deterioration despite the continued presence of intracellular MP/NP indicates that cellular damage is driven primarily by persistent extracellular MP/NP exposure rather than intracellularly retained particles.

### Global lipid remodeling induced by continuous MP/NP exposure and recovery despite intracellular MP/NP retention

To identify the specific lipid metabolites contributing to MP/NP-induced metabolic dysfunction, we performed lipidomic profiling and compared intracellular lipid compositions across different MP/NP exposure durations (T24 vs. T72) as well as between continuous exposure and recovery after extracellular MP/NP removal (T24 vs. T24-W72). Continuous MP/NP exposure resulted in extensive metabolic alterations (Fig. 3A), which is dramatically different when extracellular MP/NP are removed (Fig. 3B). Among these differentially presented lipids, AA emerged as a unique lipid, as it is significantly increased following prolonged MP/NP exposure, whereas its level decreased significantly after MP/NP removal (Fig. 3C). Comparative analysis of metabolic profiles across exposure (T24, T72) and removal (T24-W24, T24-W72) conditions further demonstrated distinct clustering patterns, indicating substantial remodeling of the cellular lipid landscape (Fig. 3D). Principal component analysis (PCA) revealed clear multidimensional separation among T24, T72 and T0 groups, while the metabolic profiles of T24-W24 and T24-W72 groups clustered closely with the T0 group (Fig. 3E), consistent with metabolic recovery after extracellular MP/NP removal. Kyoto Encyclopedia of Genes and Genomes (KEGG) pathway analysis identified unsaturated fatty acid biosynthesis as a major signaling axis significantly affected by MP/NP exposure (Fig. 3F). Heatmap exhibited significant increases in signaling and membrane-associated lipids, including AA, with continuous MP/NP exposure, whereas extracellular MP/NP removal restored these metabolites to levels comparable to untreated controls (Fig. 3G). Notably, triacylglycerol (TAG) abundance remained unchanged under both exposure and removal conditions (Fig. S8).

**Fig. 3.**
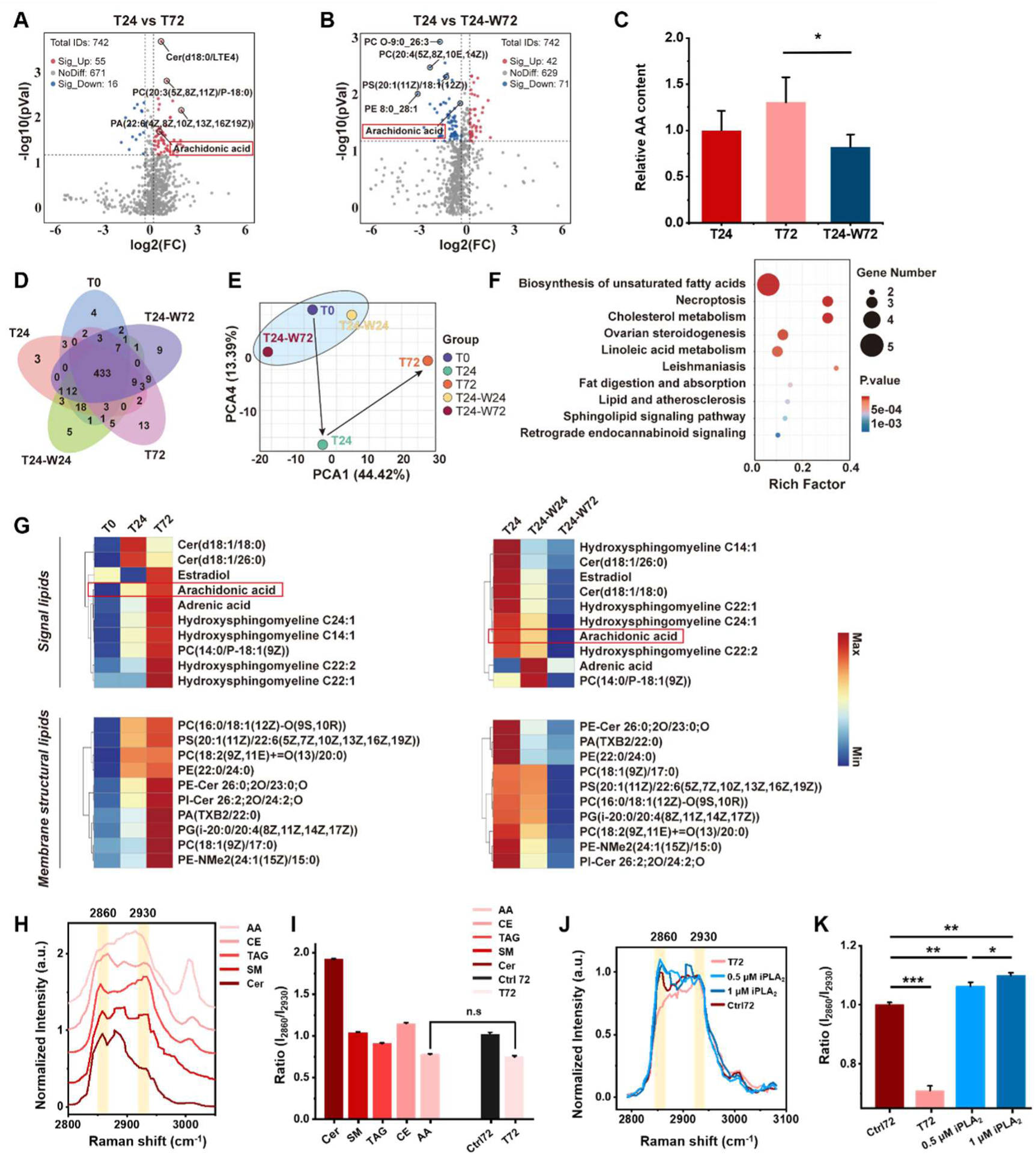
Cells continuously treating MP/NP increased the production of AA with while it can be recovered after the withdrawal of MP/NP. (**A** and **B**) Volcano plot of differential lipids contents, satisfying the criteria of log 2 (fold-change) value >1 or <-1 and P<0.05. Significantly upregulated or downregulated lipids are represented as red or blue dots, and black dots indicate nonsignificant results. The positions of AA are highlighted. (**C**) Relative AA content in cells derived from LC-MS-based analysis. (**D**) Venn diagram of differential lipids. (**E**) PCA of lipids metabolites. The blue elliptical regions represent the confidence intervals for the similarity within groups, while the arrows highlight the multivariate differences among the groups. (**F**) The most significant KEGG enrichment analyses of increased lipids metabolites in MP/NP treated groups cells. (**G**) Heatmap of representative signal lipids and membrane structural lipids among different groups cells. Red indicates upregulation, blue indicates downregulation. The positions of AA are highlighted. (**H**) SRS spectra of standard AA, CE, TAG, SM and Cer at 2800-3100 cm^-1^. (**I**) The ratio of I_2860_/I_2930_ from the SRS spectra among AA, CE, TAG, SM, Cer and LD from Ctrl72, T72 groups. (**J**) SRS spectra at 2800-3100 cm^-1^ of LDs extracted from MP/NP-treated cells in T72, 0.5 μM iPLA_2_, 1 μM iPLA_2_ and no treated control Ctrl72 groups. The characteristic peaks are labeled and highlighted. (**K**) The ratio of I_2860_/I_2930_ from the SRS spectra among T72, 0.5 μM iPLA_2_, 1 μM iPLA_2_ and no treated control Ctrl72 groups. Values are mean ± SD (n=3). * indicate P < 0.05. ** indicate P < 0.01. *** indicate P < 0.001. n.s, no significance. LC-MS, liquid chromatograph mass spectrometer; PCA, principal component analysis; KEGG, kyoto encyclopedia of genes and genomes; CE, cholesteryl ester; TAG, triacylglycerol; SM, sphingomyeline; Cer, ceramide. PLA_2_, phospholipaseA2.

### MP/NP induces spatially confined and transient alterations in single LD composition

To elucidate the underlying source of the reduced 2860/2930 ratio in LDs, we compared SRS spectra of common lipid species in LDs (triacylglycerol and cholesteryl ester) and lipid species identified from lipidomic profiling that exhibited a similar reversible trend (AA, hydroxysphingomyelin, and ceramide). Because hydroxysphingomyelin is not commercially available, sphingomyelin was used as a substitute for spectral measurements. Distinctive SRS spectra were observed for each lipid species (Fig. 3H). Notably, only AA showed a spectral profile closely resembling that of LDs from MP/NP-treated cells. Quantitative analysis of the 2860/2930 ratio across all spectra further confirmed a significant reduction in the AA spectrum (Fig. 3I), suggesting the enrichment of AA in the MP/NP treated LDs. These observations suggest that substantial intracellular AA release occurs during continuous MP/NP exposure and that AA is subsequently incorporated into LDs, giving rise to the observed LD spectral alterations. Because AA is primarily released by phospholipase A2 (PLA_2_) [36, 37], we treated MP/NP-exposed cells (72 h) with a PLA_2_ inhibitor (iPLA_2_). Inhibition of PLA_2_ resulted in a significant increase in the LD 2860/2930 ratio compared with the T72 group (Fig. 3J, K), further supporting the potential role for AA in mediating LD compositional changes.

We next investigated whether AA-associated LD alterations represent a localized metabolic response triggered during MP/NP entry. In cells from the T24 group, we identified LDs located proximal to MP/NP and LDs located distal from MP/NP within the same cells (Fig.4A, B). Hyperspectral SRS analysis revealed that the 2860 cm⁻¹ peak intensity in MP/NP-proximal LDs (LD1-3) was consistently lower than that in MP/NP-distal LDs (LD4-6, Fig. 4C). Statistical analysis confirmed that the 2860/2930 ratio was significantly reduced in MP/NP-proximal LDs compared with MP/NP-distal LDs (Fig. 4D). Importantly, this spatial heterogeneity in LD composition was no longer observed in cells containing intracellular MP/NP after extracellular MP/NP removal (T24-W72) (Fig. 4E, F), where proximal and distal LDs exhibited comparable spectral profiles (Fig. 4G) and 2860/2930 ratio (Fig. 4H). This finding indicates that localized LD compositional alterations are transient and occur specifically during active MP/NP uptake, rather than as a consequence of long-term intracellular MP/NP retention. Altogether, these observations suggest that freshly internalized MP/NP may locally stimulate AA release, potentially as part of a stress response during MP/NP uptake, thereby inducing spatially confined and transient lipid metabolic dysregulation (Fig. 4I).

**Fig. 4.**
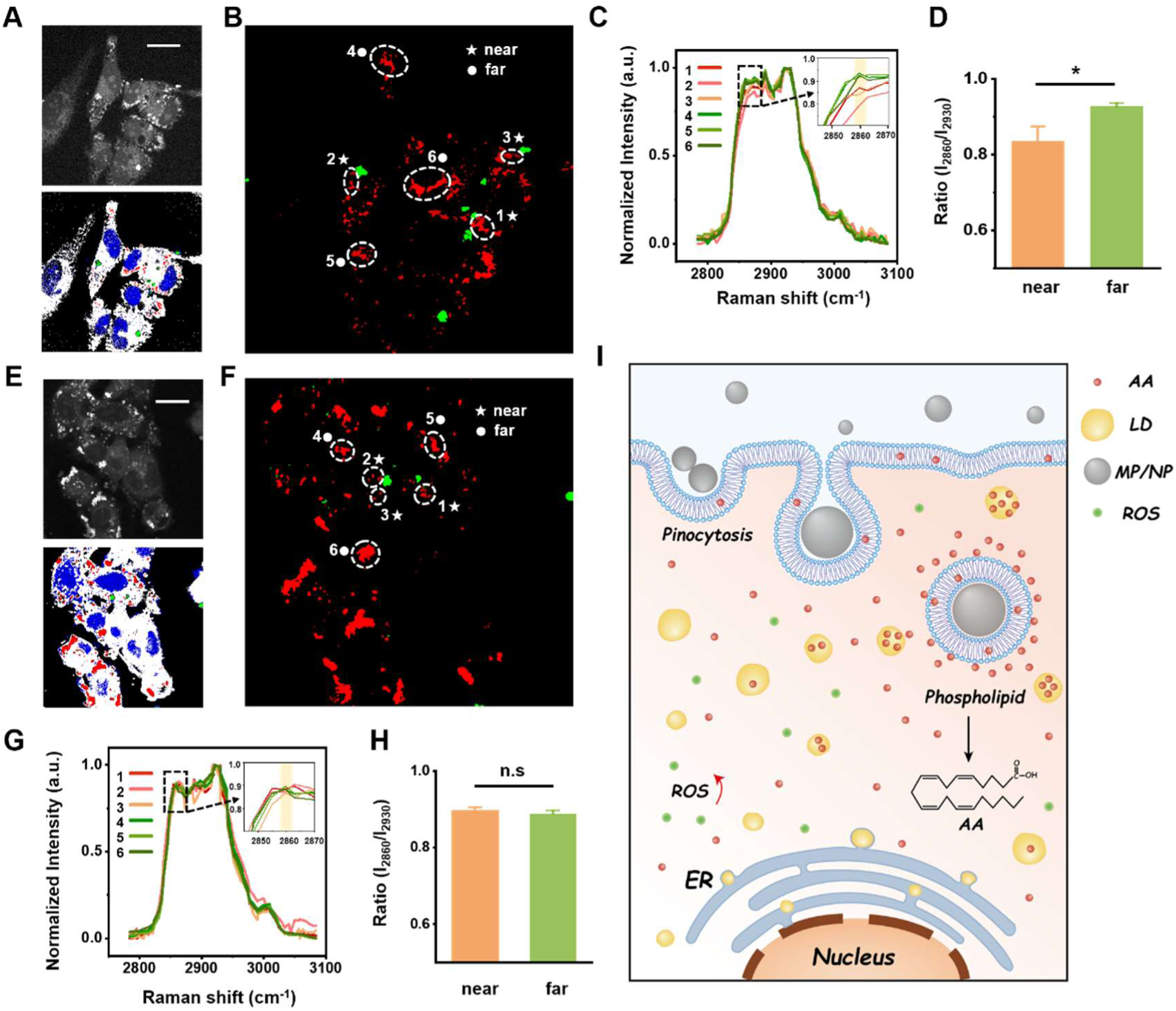
The composition of LDs shows spatial differences. (**A**) Represent SRS images at 2913 cm^-1^ and corresponding merged images from phasor analysis of T24 group cells. Red, blue, white, and green represent lipid, nucleic acid, protein and MP/NP respectively. (**B**) Corresponding spatial distribution of LDs (red) and MP/NP (green). The region within the white dotted line is the LDs to be analyzed. The stars represent the regions near to the MP/NP, while the dots represent the regions far from the MP/NP. (**C**) Spectra of LDs extracted from the regions of figure B at 2800-3100 cm^-1^. Zoom in figure shows the spectra at 2845-2870 cm^-1^. The orange or green series lines represent near to or far from the MP/NP. (**D**) The ratio of I_2860_/I_2930_ from the SRS spectra among LDs near to the MP/NP or far from the MP/NP groups. (**E**) Represent SRS images at 2913 cm^-1^ and corresponding merged images from phasor analysis of T24-W72 group cells. (**F**) Corresponding spatial distribution of LDs (red) and MP/NP (green). The region within the white dotted line is the LDs to be analyzed. The stars represent the regions near to the MP/NP, while the dots represent the regions far from the MP/NP. (**G**) Spectra of LDs extracted from the regions of figure F at 2800-3100 cm^-1^. Zoom in figure shows the spectra at 2845-2870 cm^-1^. The orange or green series lines represent near to or far from the MP/NP. (**H**) The ratio of I_2860_/I_2930_ from the SRS spectra among LDs near to the MP/NP or far from the MP/NP groups. (**I**) Schematic diagram of MP/NP ingestion on lipid metabolism mechanisms. Scale bar, 10 μm. Values are mean ± SD (n=3). * indicate P < 0.05. n.s, no significance.

### Conserved across diverse cell types and particles

To determine whether the observed phenomenon is a general cellular response, we extended the study to test polystyrene (PS), a widely used MP/NP model (Fig. S9A). The SRS spectrum of PS MP/NP showed a prominent peak at 3055 cm^-1^, allowing for the bond-selective imaging of PS MP/NP and LDs within cells (Fig. S9B, C). Consistent with the findings for clinically derived MP/NP, a significant reduction in the 2860/2930 ratio was observed exclusively in LDs proximal to PS MP/NP, while distal LDs remained metabolically stable (Fig. S9D-J). This spatial heterogeneity in LD composition disappeared after the removal of extracellular PS MP/NP (Fig. S9K-M). Functionally, the toxicity and cellular damage associated with PS MP/NP were only apparent when particles were continuously present in the extracellular medium, and the cells began to recover once extracellular PS MP/NP were removed (Fig. S9N, O). Furthermore, to test whether this phenomenon is cell-type specific, we evaluated the effects in other cell types, including HUVECs (human endothelial cells) and A549 (human lung cancer cells). Both cell types exhibited the same reversible reduction in the 2860/2930 ratio upon MP/NP exposure (Fig. S10). Together, these results suggest that localized metabolic remodeling of LDs and transient cytotoxicity are likely conserved across diverse cell types and MP/NPs compositions.

### Phospholipid coating attenuates uptake-associated MP/NP cytotoxicity

Motivated by these findings, we next tested whether surface-mediated interactions during internalization drive MP/NP-induced cytotoxicity, but not intracellular particle retention. To test this hypothesis, we coated MP/NP with a phospholipid layer to mimic a more biocompatible surface and potentially mitigate harmful interactions during cellular uptake. The integrity and completeness of the phospholipid coating were validated by incorporating the fluorescent probe 3,3’-dioctadecyloxacarbocyanine perchlorate (DiO) into an artificial phospholipid membrane composed of 1,2-Dioleoyl-sn-glycero-3-phosphocholine (DOPC) (Fig. 5A). Merged fluorescence-brightfield images confirmed uniform coating of DOPC-DiO on MP/NP (Fig. 5B). Surface chemical composition analysis using energy-dispersive X-ray spectroscopy (EDS) further supported successful coating. Both natural MP/NP and DOPC-DiO MP/NP exhibited strong carbon (C) and oxygen (O) signals, whereas a distinct phosphorus (P) signal (derived from DOPC) was detected exclusively on DOPC-DiO MP/NP (Fig. 5C, D). These results collectively validate the successful phospholipid coating of MP/NP.

**Fig. 5.**
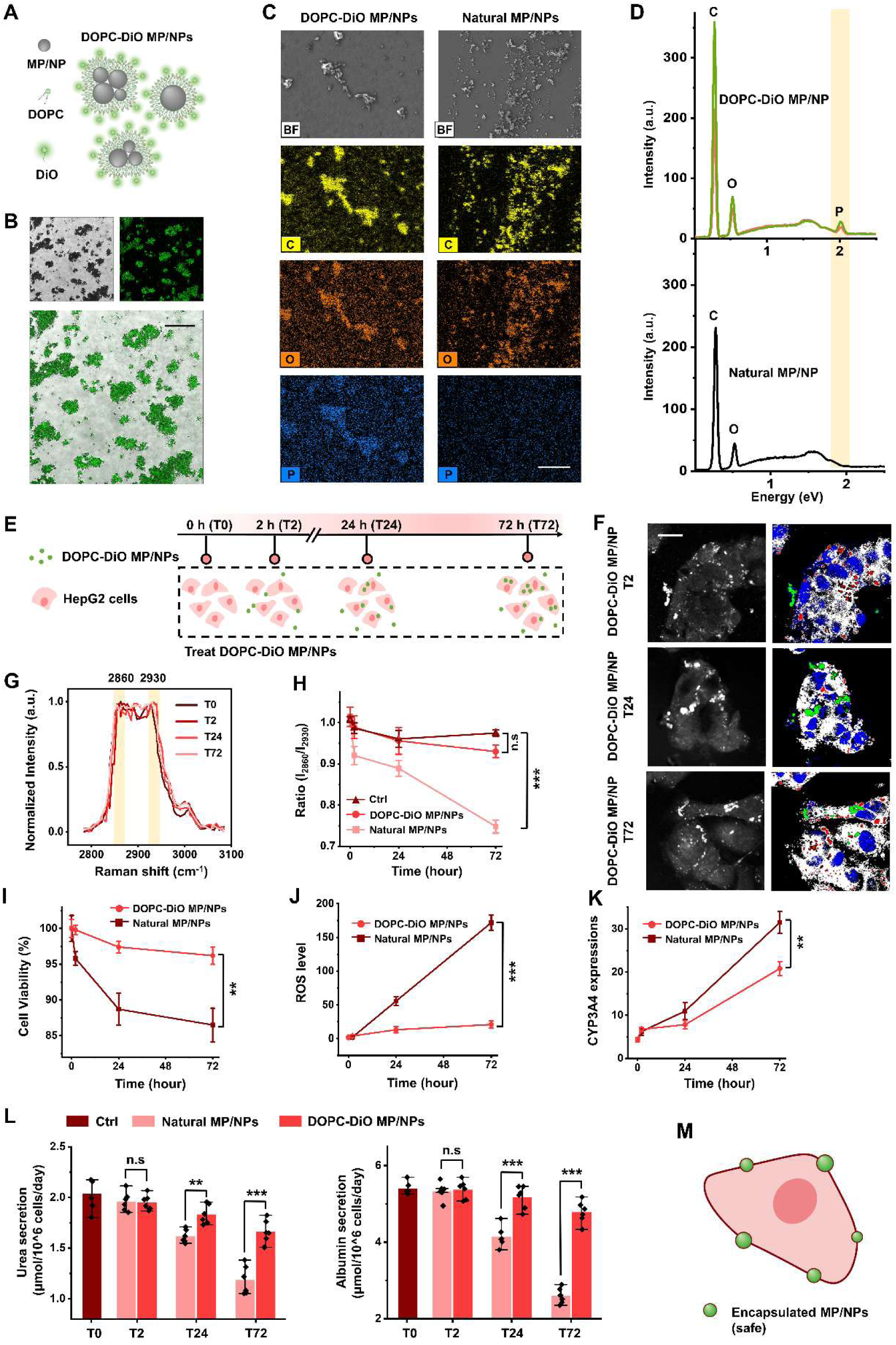
DOPC-DiO MP/NP reduced cytotoxicity effectively. (**A**) Diagram of preparing DOPC-DiO MP/NP. (**B**) Fluorescence (up right) and bright field (up left) merge images (bottom) of DOPC-DiO MP/NP. Scale bar, 200 μm. (**C**). Elemental mapping revealed that DOPC-DiO MP/NP surfaces are mainly C (yellow), O (orange) and P (blue) elements, while natural MP/NP, surfaces are mainly C (yellow) and O (orange) elements. Scale bar, 50 μm. (**D**) Corresponding EDX spectra of DOPC-DiO MP/NP and natural MP/NP surfaces. The green and yellow lines represent different detection sites of the same sample. The characteristic peaks are labeled and highlighted. (**E**) Schematic diagram of the experimental timeline for treating cells with DOPC-DiO MP/NP. (**F**) Representative SRS images at 2913 cm^-1^ and corresponding merged images from phasor analysis. Red, blue, white, and green represent lipid, nucleic acid, protein and MP/NP respectively. Scale bar, 10 μm. (**G**) SRS spectra at 2800-3100 cm^-1^ of LDs extracted from DOPC-DiO MP/NP-treated cells in T0, T2, T24 and T72 groups. The characteristic peaks are labeled and highlighted. (**H**) The ratio of I_2860_/I_2930_ from the SRS spectra comparing DOPC-DiO MP/NP-treated cells (red line), natural MP/NP-treated cells (light red line) and control cells (brown line). (**I**) Cell viability evaluated by CCK-8 assay in DOPC-DiO MP/NP-treated and natural MP/NP-treated groups. (**J**) ROS levels in DOPC-DiO MP/NP-treated and natural MP/NP-treated groups. (**K**) Quantification of CYP3A4 expressions. (**L**) Urea and albumin secretion were examined by ELISA kit. (**M**) Diagram of the cell continuously treat with encapsulated MP/NP. Values are mean ± SD (n=3). ** indicate P < 0.01. *** indicate P < 0.001. n.s, no significance. EDX, energy-dispersive X-ray spectroscopy; C, carbon; O, oxygen; P, phosphorus.

The DOPC-DiO MP/NP were then used to treat cells to assess their cytotoxicity (Fig. 5E). Hyperspectral SRS images show similar MP/NP uptake and distribution after phospholipid coating (Fig. 5F). Interestingly, SRS spectra of LDs revealed that LDs from DOPC-DiO MP/NP-treated cells showed spectral profiles similar to those of time-matched untreated controls (Fig. 5G). Quantitative analysis further showed that 2860/2930 ratio maintained similar to the untreated control cells in the DOPC-DiO MP/NP group, showing significantly different spectral profiles compared to the uncoated MP/NP (Fig. 5H). Consistent with the spectral findings, CCK-8 assays showed that the viability is recovered significantly in DOPC-DiO MP/NP-treated cells compared to the uncoated MP/NP (Fig. 5I). Intracellular ROS levels were significantly reduced following DOPC-DiO MP/NP exposure compared to uncoated MP/NP exposure (Fig. 5J, Fig. S11A). CYP3A4 expression levels were significantly reduced in DOPC-DiO MP/NP-treated cells compared to the uncoated MP/NP (Fig. 5K, Fig. S11B). The secretion function of hepatocytes remained intact upon continuous DOPC-DiO MP/NP exposures, measured by significantly higher urea and albumin levels in the medium (Fig. 5L). These results suggest that phospholipid coating has effectively shielded the damaging effect of MP/NP, protecting cells from MP/NP attack on the membrane (Fig. 5M). Together, these findings support the hypothesis that surface chemistry and the cellular internalization process are major contributors of MP/NP-induced cellular damage.

## Discussion

Although the adverse biological effects of MP/NP are increasingly recognized, the specific mechanisms by which they induce stress during internalization remain unclear [38–40]. This gap stems in part from the technical challenge of tracking naturally derived MP/NP within cells with sufficient spatial resolution and chemical specificity. Commonly used detection approaches, such as thermal pyrolysis gas chromatography-mass spectrometry, fluorescence labeling, and electron microscopy, are challenged by sample destruction, complex preparation procedures, and limited quantitative capability for dynamic tracking [41–43]. Recent discussions have further highlighted widespread methodological concerns in MP/NP research, highlighting an urgent need for detection strategies that are both chemically specific and quantitative [44]. Here, we address this need using a vibrational spectroscopic imaging approach (SRS) that enables label-free, chemically specific identification of MP/NP in complex cellular environments and direct tracking of their interactions at the subcellular level. Using this strategy, we found a previously unrecognized mechanism of MP/NP-induced cellular damage at the single-cell level, providing mechanistic insight that may inform future strategies for mitigating the human health risks associated with MP/NP exposure.

By visualizing the spatiotemporal dynamics of MP/NP uptake using SRS microscopy, we observed a dynamic internalization process in which MP/NP progress from membrane association to cytoplasmic dispersion and aggregation. This direct visualization clarifies the intracellular fate of MP/NP and establishes a way for understanding their downstream biological consequences. Notably, hyperspectral SRS analysis of LDs revealed that continuous MP/NP exposure induces significant alterations in LD composition, whereas removal of extracellular MP/NP leads to substantial metabolic recovery. Importantly, despite these compositional changes, neither LD size nor LD area fraction differed significantly across different MP/NP exposure conditions (Fig. S12), in contrast to previous reports of MP/NP-induced LD accumulation in zebrafish [31]. This discrepancy likely reflects differences in biological scale and mechanism, as our study captures the direct, primary effects of cellular MP/NP internalization, whereas in vivo models reflect secondary systemic responses that promote LD accumulation under prolonged organismal exposure.

Mechanistically, we identified AA, a key inflammatory mediator and cancer-associated lipid species [45–47], as a pivotal regulator of MP/NP-induced metabolic dysregulation. The strong correlation between the Raman spectral signature of AA standards and those of MP/NP-affected LDs, together with the restoration of the 2860/2930 cm^-1^ ratio following PLA_2_ inhibition, supports a potential role for PLA_2_ in mediating AA enrichment in LDs during MP/NP exposure. Because this process typically occurs at the plasma membrane, it is possible that local AA release near freshly internalized MP/NP arises from partial membrane disruption during MP/NP internalization, depending on MP/NP surface chemistry. Future validation of this model will require spatially resolved quantification of PLA_2_ activity at the plasma membrane proximal to MP/NP internalization. Indeed, MP/NP can interact with membranes through multiple mechanisms, including surface adsorption that masks membrane receptors [48], mechanical deformation that triggers stress responses, physical penetration of the lipid bilayer leading to content leakage [49, 50], pore formation that increases membrane permeability [51], and ROS-induced lipid peroxidation that reduces membrane fluidity [52]. Strikingly, phospholipid coating, which is designed to mimic a more biocompatible surface chemistry for cellular uptake, effectively abrogated localized AA release near internalized MP/NP and eliminated associated cytotoxicity, presenting a safe entry mode of internalization.

Importantly, the consistent reduction in the 2860/2930 ratio across diverse cell types strongly suggests that the AA-mediated pathway is a generalized reaction to MP/NP-lipid interactions. Furthermore, the observation that chemically distinct particles (PMMA and PS) exhibit the same localized LD compositional alterations indicates that, for these chemically inert cores, the physical mechanics of internalization are the critical driver of damage. The transient nature of this stress response was further validated by the dynamics of CYP3A4, a key hepatic detoxification enzyme. The upregulation of CYP3A4 during MP/NP exposure suggests a cellular defense response to these foreign particles [34, 35], likely mediated by xenobiotic-sensing pathways such as PXR [53, 54]. Notably, the subsequent decline in CYP3A4 expression following MP/NP washout indicates a return to baseline function, further demonstrating MP/NP as a reversible stress. Together, these results provide direct evidence for the chemically inert nature of MP/NP cores while revealing how and at which stage cellular damage arises. More broadly, our findings prompt a reassessment of MP/NP health risks by highlighting uptake-associated, localized and transient metabolic stress as a critical source of toxicity, thereby informing future strategies for evaluating and mitigating MP/NP exposures.

## Materials & Methods

### Cell lines, cell culture, and chemicals

Human hepatocellular carcinoma cells HepG2, human umbilical vein endothelial cells HUVEC, and human lung cancer cells A549 cells were obtained from the American Type Culture Collection (ATCC). HepG2 cells were cultured in MEM (MeilunBio, MA0217) with 15% fetal bovine serum (Procell, 164210) and incubated at 37℃ and 5% CO_2_. HUVEC and A549 cells were cultured in DMEM (Gibco, Cat #11995065) with 10% fetal bovine serum (Procell, 164210) and incubated at 37℃ and 5% CO_2_. For exposure experiments, cells were seeded at 1 × 10^5^cells/mL and exposed to 10 μg/mL MP/NP for 2 h, 24 h, or 72 h to be comparability with human exposure level (12 μg/ml in the blood) (*33*). Sphingomyelin and ceramide were purchased from Aladdin (Cat# S130561 and N130628).

### MP/NP preparation and characterization

MP/NP were generated from commercially available medical-grade bone cement (Landmover Medical, Cat# PALACOS-H). As previously described (*55, 56*), the bulk cement was sectioned and subjected to primary grinding using a pulverizer (Taist, FW135). The resulting fragments were sieved through a 50-μm mesh to isolate particles smaller than 50 μm. Particles passing the sieve were further refined using a planetary ball mill. This secondary milling step was followed by vacuum filtration through a 1-μm membrane filter to isolate MP/NP. After natural drying, the final processed submicron MP/NP were obtained for cell exposure and label-free SRS imaging. PS MP/NP were purchased from Haian Zhichuan Battery Material Technology (Cat# ps-0001).

Particle size and zeta potential were determined by Zetasizer Nano-ZS. Nanoparticles were diluted in purified water to 0.01 mg/mL and measured at 37°C. Data were collected from three replicates. For fluorescence imaging, MP/NP were coated with fluorescence-labeled phospholipids. Briefly, 1,2-dioleoyl-sn-glycero-3-phosphocholine (DOPC, D130438, Aladdin) was mixed with the fluorescent dye DiO (C1038, Beyotime) to prepare a labeled phospholipid solution. MP/NP were dispersed in this solution and sonicated to facilitate phospholipid encapsulation, yielding a stable suspension of fluorescently coated MP/NP.

### SRS microscopy

A home-built picosecond SRS microscope (*57, 58*) was used (Fig.S1). A picosecond laser (picoEmerald) provided synchronized pump (tunable wavelength 700–990 nm and 80 MHz repetition rate) and Stokes beams (fixed wavelength at 1031 nm and 2-ps pulse width). The Stokes beam was intensity-modulated with an electro-optic modulator at 20 MHz. Both beams were directed into a laser-scanning microscope (BX51WI, Olympus) equipped with a 60X, 1.2 NA water-immersion objective (Olympus). The SRS signal was detected using a photodiode and extracted by a lock-in amplifier. Images were acquired with a pixel dwell time of 10 µs.

### Spectral phasor analysis

The SRS image data set (2800-3100 cm^-1^ with a 4 cm^-1^ step size) was imported into FIJI (Version 1.54f), and an average intensity projection was generated. The spectral phasor analysis was performed as described previously (*59*). Segmentation of the phasor plot was performed manually using regions of interest (ROIs) to create images of discrete cellular locations. The corresponding average spectra for each ROI were plotted using OriginPro (2018C).

### SEM and EDS characterization

A SEM system (SU-6600, Hitachi, Japan) was used for SEM analysis as previously described (*60*). Samples were sputter-coated with gold-platinum for 4-5 min using a sputter coater (Ion Sputter E-1045, Hitachi, Tokyo, Japan). For EDS analysis of modified fiber-grid scaffolds, samples were placed on the specimen stage and securely fixed. Elemental distribution, including nitrogen and silicon, within the ROIs were analyzed using the mapping mode.

### UPLC–MS/MS

A 200 μL sample was mixed with 400 μL ice-cold methyl tert-butyl ether (MTBE) and 80 μL ice-cold methanol. After centrifugation at 3,000 rpm for 15 min, 200 μL of the supernatant was transferred to another tube. The supernatant was freeze-dried, reconstituted with 200 μL dichloromethane/methanol (1:1, v/v), and centrifuged again at 3000 rpm for 15 min. The final supernatant was transferred to a fresh vial for UPLC-HRMS analysis. A quality control sample was prepared by mixing equal aliquots of sample supernatants.

Lipids were analyzed using a Q Exactive mass spectrometer (Thermo Fisher Scientific) with a CSH C18 column (1.7 μm, 2.1 × 100 mm; Waters, USA). The elution gradient was as follows: 0−2 min, 40%−43% solvent B (10 mM ammonia formate, 0.1% formic acid, 90% isopropyl alcohol, 10% acetonitrile); 2−2.1 min, 43%−50% solvent B; 2.1−7 min, 50%−54% solvent B; 7−7.1 min, 54%−70% solvent B; 7.1−13 min, 70%−99% solvent B; flow rate at 0.35 mL/min.

Mass spectrometric settings for positive/negative ionization modes were: spray voltage 3.8/–3.2 kV; auxiliary gas heater 350 °C; capillary temperature 320 °C. Full-scan range was 200–2000 m/z with 70,000 resolution; AGC target for MS acquisitions was set to 3e6 with a maximum injection time of 100 ms. The top three precursors were selected for MS fragmentation with a maximum ion injection time of 50 ms with 17,500 resolution, and AGC target set to 1e5. Stepped normalized collision energies were 15, 30, and 45 eV. LipidSearch 4.1 SP2 software (Thermo Fisher, USA) was used for lipid identification and quantitation.

### CCK-8 assay

HepG2 cells were seeded in 96-well plates at 103 cells/well and allowed to adhere overnight. At each time point, the medium was carefully aspirated and replaced with 100 µL fresh serum-containing medium. Then, 10 µL CCK-8 reagent (Solarbio, Cat# CA1210-500T) was added to each well. Plates were incubated at 37 °C in a humidified 5% CO₂ for 2 h, protected from light. Absorbance at 450 nm was measured with a microplate reader (TUOHE, Cat# TMR-100). Wells containing medium and CCK-8 reagent without cells served as blanks.

### Cell viability assays

Cells were washed three times with PBS, then stained using LIVE/DEAD reagents (Key-GEN BioTECH Co., Ltd, China) following the manufacturer’s instructions. Calcein-AM and propidium iodide (PI) were diluted in PBS to 2 μM and 8 μM, respectively. After incubation with the Calcein-AM/PI mixture for 30 min in the dark, samples were washed three times with PBS and imaged using a confocal laser scanning microscope (OLYMPUS FV3000). Green (ex 495/em 519 nm) and red channels (ex 555/em 565 nm) represented live and dead cells, respectively.

### ROS Assay

ROS generation was assessed using a ROS Assay Kit (Beyotime, Cat# S0035S). Cells were stained with 10 μM DCFH-DA at 37 °C for 30 min and imaged using a confocal laser scanning microscope (OLYMPUS FV3000). Each experiment included technical replicates and was repeated independently at least three times (n ≥ 3).

### ELISA assays

Secreted proteins were quantified by ELISA. Cell culture supernatants were collected and centrifuged at 1,000 rpm for 20 min to remove debris. Albumin and urea concentrations were measured using human albumin (ab108788, Abcam, USA) and human urea (ab83362, Abcam) ELISA kits, respectively, following the manufacturer’s protocols.

### Immunohistochemical staining

Cells were fixed in 4% paraformaldehyde for 10 min and permeabilized with 0.1% Triton X-100 for 10 min. Samples were blocked in PBS containing 5% serum from the same species as the secondary antibody for 30 min, then incubated overnight at 4 °C with CYP3A4 primary antibody pre-conjugated with a fluorescent secondary antibody (CL488-67110, Proteintech). After PBS washes, samples were stained with DAPI (Beyotime, Cat# C1002) for 5 min in the dark. Imaging was performed using a confocal laser scanning microscope (OLYMPUS FV3000). The primary and secondary antibodies used are listed.

### Inhibitor experiment

Cells were pre-incubated with 20 μM Chlorpromazine (MedChemExpress, Cat# 50-53-3) or 40 μM EIPA (MedChemExpress, Cat# L593754) for 60 min at 37 °C. Then, MP/NP were added to the medium for 24h. After washing with ice-cold PBS, cells were fixed with 4% paraformaldehyde and imaged.

### Statistical analysis

All data are presented as mean ± standard deviation (SD). Statistical analysis was performed using GraphPad Prism v.8. Comparisons between two groups were made using Student’s t-test. Multiple-group comparisons were performed using one-way ANOVA followed by Tukey’s post hoc test. A p-value < 0.05 was considered statistically significant.

## Acknowledgments

We thank Prof. Xinhua Yao and Dr. Lixin Tian at Zhejiang University for providing access to the planetary ball mill, and Xingxin Liu for assistance with SRS system maintenance.

## Funding

This work was supported by National Natural Science Foundation of China (W2532057, 82372011), Zhejiang Provincial Natural Science Foundation of China (LZ25H180001), and Fundamental Research Funds for the Central Universities (226-2025-00034).

## Competing interests

The authors declare they have no competing interest.

## Data and materials availability

All data needed to evaluate the conclusion in the paper are present in the paper and/or the Supplementary Materials.

## Notes

### Competing Interest Statement

The authors have declared no competing interest.

