## Supplementary Materials for "Clinically derived micro- and nanoplastics uptake drives spatiotemporally confined metabolic stress revealed by bond-selective imaging"

**This PDF file includes:**

Figs. S1 to S12

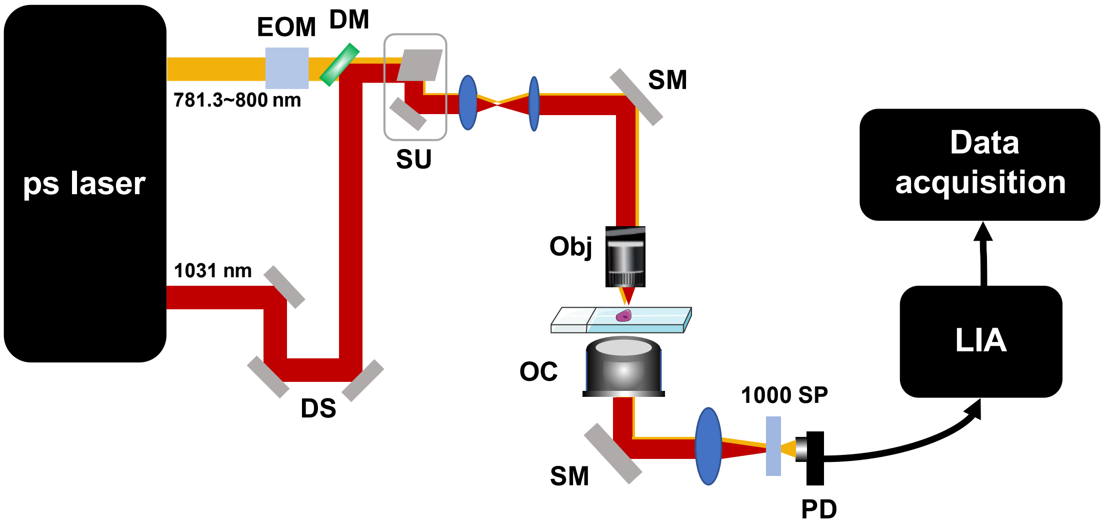


**Fig. S1. SRS microscopy.** EOM, electro-optic modulator; DM, dichroic mirror; DS, delay stage; SU, scan unit; SM, silver mirror; Obj, objective; OC, oil condenser; SP, short pass; PD, photodiode; LIA, lock-in amplifier.

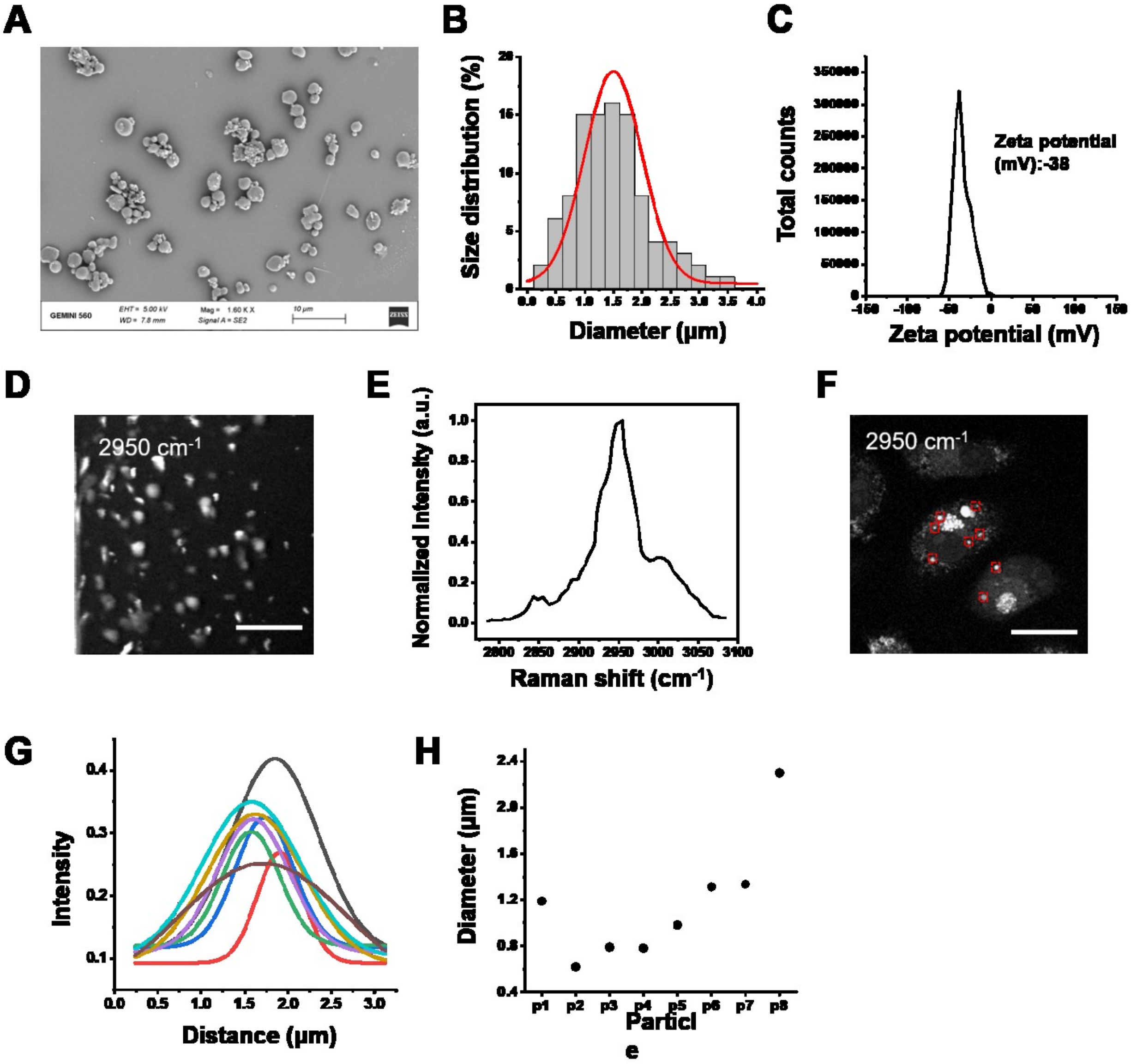


**Fig. S2. Characterization of MP/NPs.** (**A**) SEM image of MP/NPs. (**B**) Size distribution of MP/NPs. (**C**) Zeta potential of MP/NPs. (**D**) Representative SRS image of MP/NPs at 2950 $cm^{-1}$. Scale bar, 10 μm. (**E**) Corresponding SRS spectra at 2800-3100 $cm^{-1}$ of the MP/NPs. (**F**) SRS image of distribution of plastics within live cells (also presented in Fig.1B). (**G**) Gaussian fitting of monodisperse plastics. (**H**) Diameters of monodisperse plastic particles.

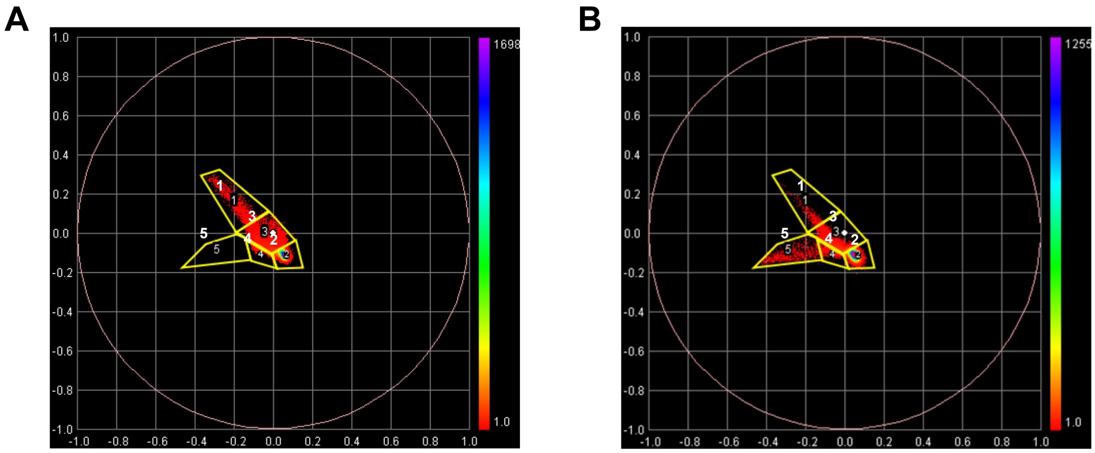


**Fig. S3. Represented phasor plot of SRS data of MP/NP uptake cells.** (**A**) Cells not treated with MP/NPs. (**B**) Cells treated with MP/NPs. ROI 1 to 5 represent lipid, background, protein, nucleic acid, and MP/NPs components respectively.

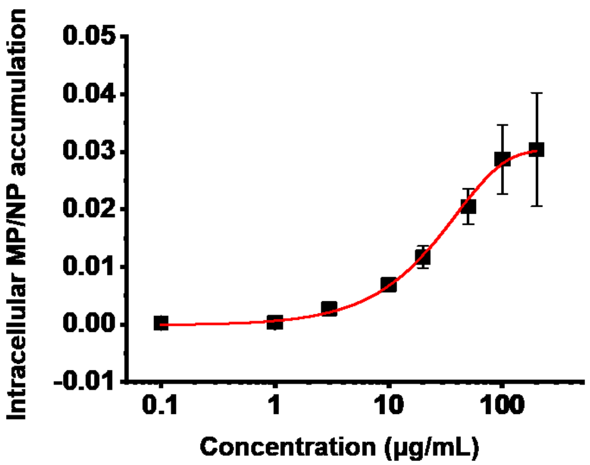


**Fig.S4. Intracellular MP/NP accumulation in T2 group cells with different MP/NP concentration.** Values are mean ± SD (n=3).

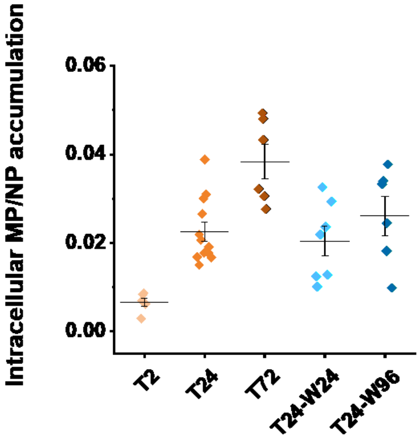

**Fig. S5. Intracellular MP/NP accumulation at different time point.** Values are mean ± SD (n=6 in T2, n=12 in T24, n=6 in T72, n=7 in T24-W24, n=6 in T24-W72).

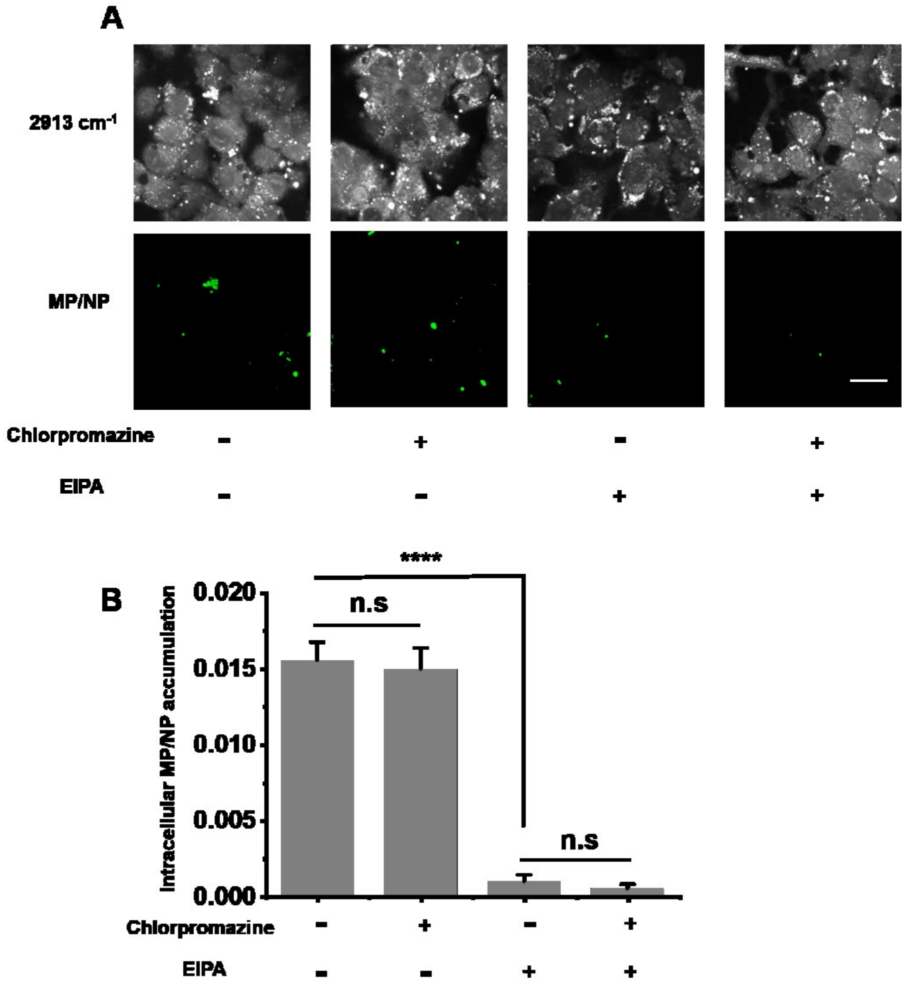


**Fig. S6. Inhibitor experiment of endocytosis and pinocytosis markers.** (**A**) Representative SRS images at 2913 $cm^{-1}$ and corresponding MP/NP images from phasor analysis. Scale bar, 10 µm. (**B**) Intracellular MP/NP accumulation statistics chart. Values are mean ± SD (n=3). **** indicate P < 0.0001. n.s, no significance.

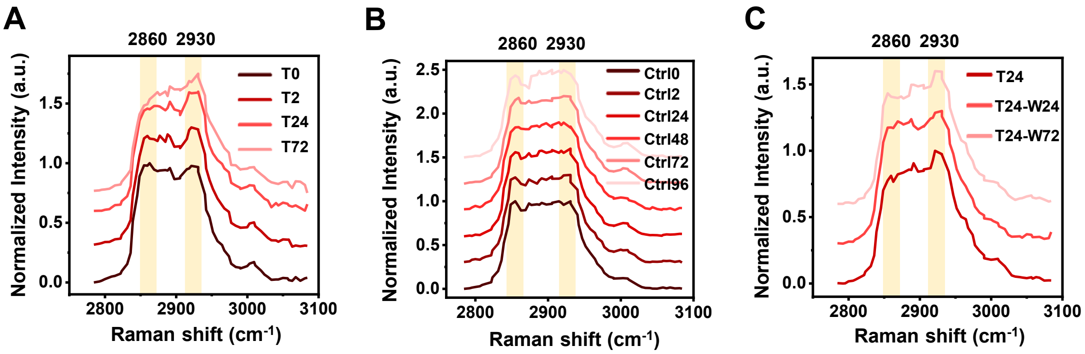


**Fig. S7.** Offset SRS spectra at 2800-3100 $cm^{-1}$ of LDs extracted from MP/NP-treated cells in T0, T2, T24 and T72 groups (**A**) and control cells in Ctrl0, Ctrl2, Ctrl24, Ctrl48, Ctrl72 and Ctrl96 groups (**B**). (**C**) Offset SRS spectra of LDs extracted from MP/NP-washout cells in T24, T24-W24 and T24-W72 groups.

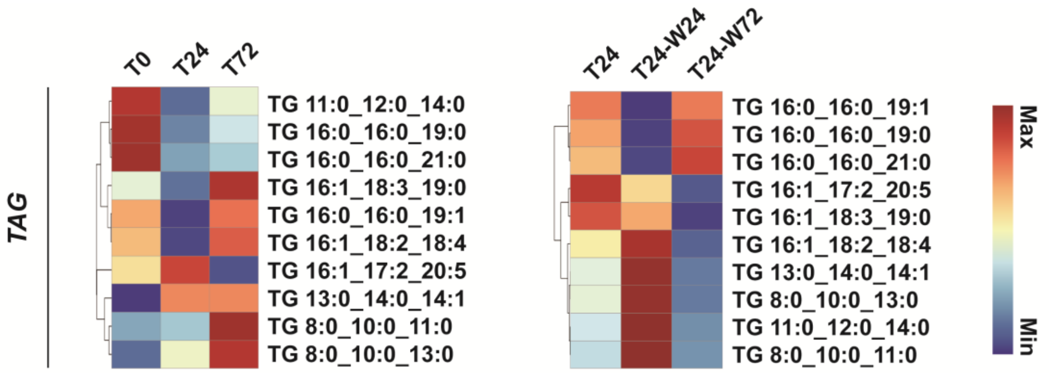


**Fig. S8. Heatmap of representative TAG among different groups cells.** The abscissa represents the sample name, and the ordinate represents the clustering results for the differentially lipids metabolites. Red indicates upregulation, and blue indicates downregulation. TAG, triacylglycerol.

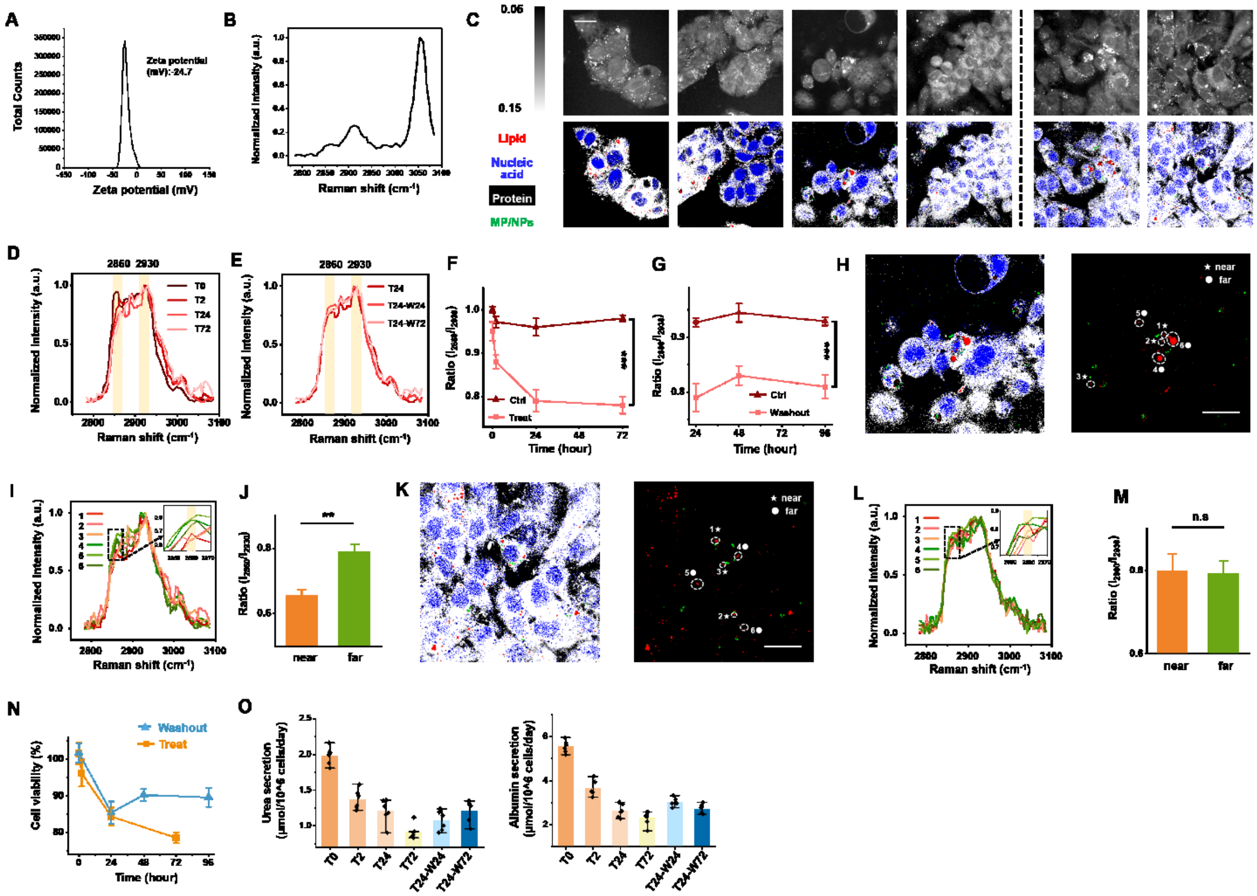


**Fig. S9. PS MP/NP-treated HepG2 cells exhibited a similar trend of LD composition, spatial differences, and functions.** (A) Zeta potential of PS MP/NP. (B) SRS spectra at 2800-3100 cm$^{-1}$ of the PS MP/NP. (C) Representative SRS images at 2913 cm$^{-1}$ and corresponding merged images from phasor analysis. Red, blue, white, and green represent lipid, nucleic acid, protein and PS MP/NP respectively. Scale bar, 10 μm. (D) SRS spectra at 2800-3100 cm$^{-1}$ of LDs extracted from PS MP/NP-treated cells in T0, T2, T24 and T72 groups. (E) SRS spectra at 2800-3100 cm$^{-1}$ of LDs extracted from PS MP/NP-treated cells in T24, T24-W24 and T24-W72 groups. (F) The ratio of $I_{2860}/I_{2930}$ from the SRS spectra comparing PS MP/NP-treated cells (red line) and control cells (brown line). (G) The ratio of $I_{2860}/I_{2930}$ from the SRS spectra comparing PS MP/NP-washout cells (red line) and control cells (brown line). (H) Represent merged images from phasor analysis (also presented in C) and corresponding spatial distribution of LDs (red) and MP/NP (green) from phasor analysis of T24 group cells. The region within the white dotted line is the LDs to be analyzed. Each area is marked with numbers in sequence. The stars represent the regions near to the MP/NP, while the dots represent the regions far from the MP/NP. Red, blue, white, and green represent lipid, nucleic acid, protein and PS MP/NP respectively. Scale bar, 10 μm. (I) Spectra of LDs at 2800-3100 cm$^{-1}$. Zoom in figure shows the spectra at 2845-2870 cm$^{-1}$. The orange series lines represent close distance to the PS MP/NP, the green series lines represent far from the PS MP/NP. The characteristic peak is highlighted. (J) The ratio of $I_{2860}/I_{2930}$ from the SRS spectra among LDs near to the PS MP/NP or far from the PS MP/NP groups. (K) Represent merged images from phasor analysis (also presented in C) and corresponding spatial distribution of LDs (red) and MP/NP (green) from phasor analysis of T24-W72 group cells. The region within the white dotted line is the LDs to be analyzed. Each area is marked with numbers in sequence. The stars represent the regions near to the MP/NP, while the dots represent the regions far from the MP/NP. Red, blue, white, and green represent lipid, nucleic acid, protein and PS MP/NP respectively. Scale bar, 10 μm. (L) Spectra of LDs at 2800-3100 $cm^{-1}$. Zoom in figure shows the spectra at 2845-2870 $cm^{-1}$. The orange series lines represent close distance to the PS MP/NP, the green series lines represent far from the PS MP/NP. The characteristic peak is highlighted. (M) The ratio of $I_{2860}/I_{2930}$ from the SRS spectra among LDs near to the PS MP/NP or far from the PS MP/NP groups. (N) Cell viability evaluated by CCK-8 assay in PS MP/NP-treat and PS MP/NP-washout groups. (O) Urea and albumin secretion were examined by ELISA kit. Values are mean ± SD (n=3). *** indicate $P < 0.001$. ** indicate $P < 0.01$. n.s, no significance.

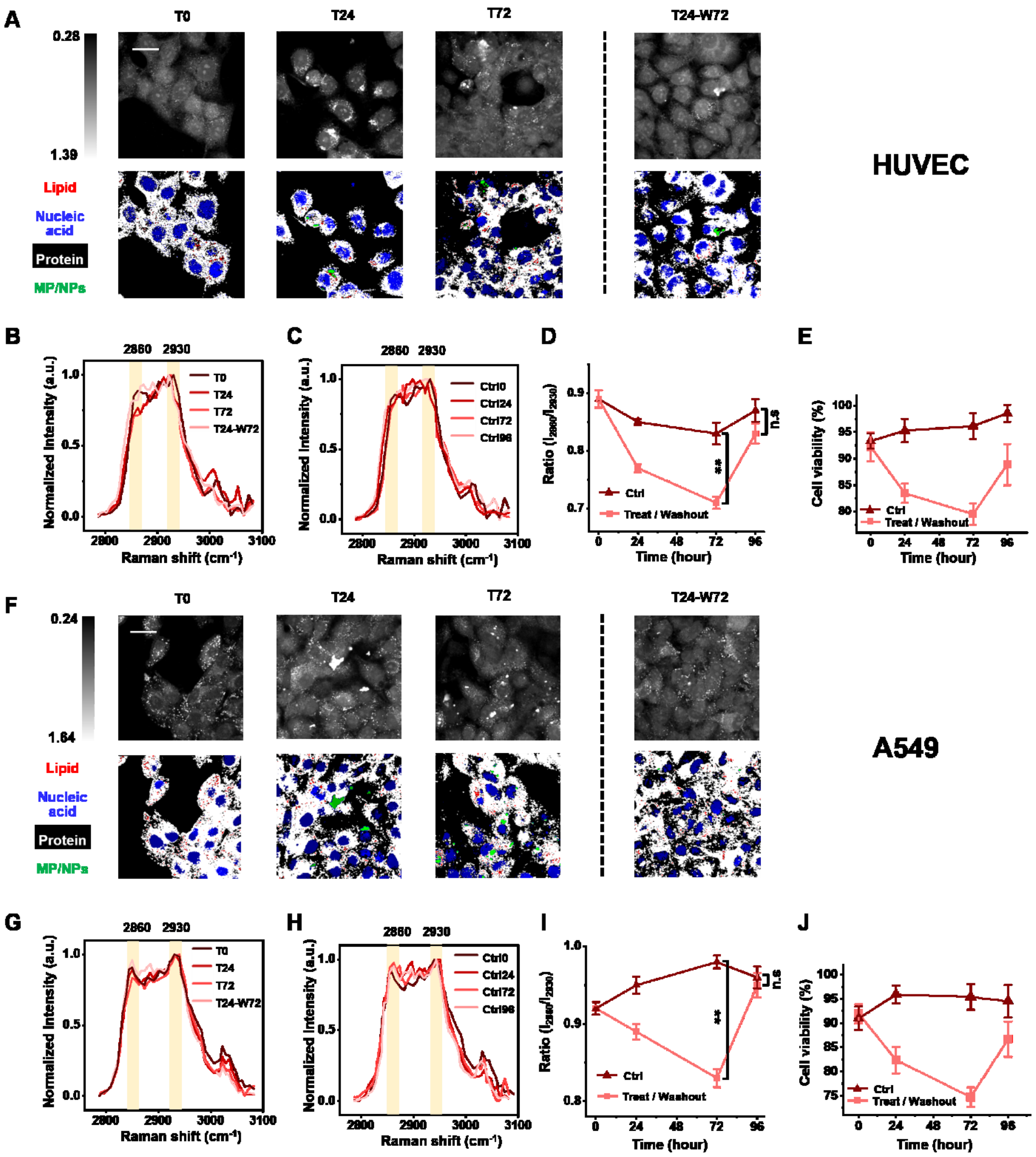


**Fig. S10. MP/NP-treated HUVEC and A549 cells exhibited similar effect.** (**A**) Representative SRS images at 2913 $cm^{-1}$ and corresponding merged images from phasor analysis in HUVEC. Red, blue, white, and green represent lipid, nucleic acid, protein and MP/NP respectively. Scale bar, 10 µm. (**B**) SRS spectra at 2800-3100 $cm^{-1}$ of LDs extracted from MP/NP-treat/washout HUVEC in T0, T24, T72 and T24-W72 groups. (**C**) SRS spectra at 2800-3100 $cm^{-1}$ of LDs extracted from control HUVEC in Ctrl0, Ctrl24, Ctrl72 and Ctrl96 groups. (**D**) The ratio of $I_{2860}/I_{2930}$ from the SRS spectra comparing MP/NP-treat/washout HUVEC (red line) and control HUVEC (brown line). The characteristic peak is highlighted. (**E**) HUVEC viability evaluated by CCK-8 assay in MP/NP-treat/washout groups. (**F**) Representative SRS images at 2913 $cm^{-1}$ and corresponding merged images from phasor analysis in A549 cells. Red, blue, white, and green represent lipid, nucleic acid, protein and MP/NP respectively. Scale bar, 10 μm. (**G**) SRS spectra at 2800-3100 $cm^{-1}$ of LDs extracted from MP/NP-treat/washout A549 cells in T0, T24, T72 and T24-W72 groups. (**H**) SRS spectra at 2800-3100 $cm^{-1}$ of LDs extracted from control A549 cells in Ctrl0, Ctrl24, Ctrl72 and Ctrl96 groups. (**I**) The ratio of $I_{2860}/I_{2930}$ from the SRS spectra comparing MP/NP-treat/washout A549 cells (red line) and control A549 cells (brown line). The characteristic peak is highlighted. (**J**) A549 cells viability evaluated by CCK-8 assay in MP/NP-treat/washout groups. The characteristic peak is highlighted. Values are mean ± SD (n=3). ** indicate P < 0.01. n.s, no significance.

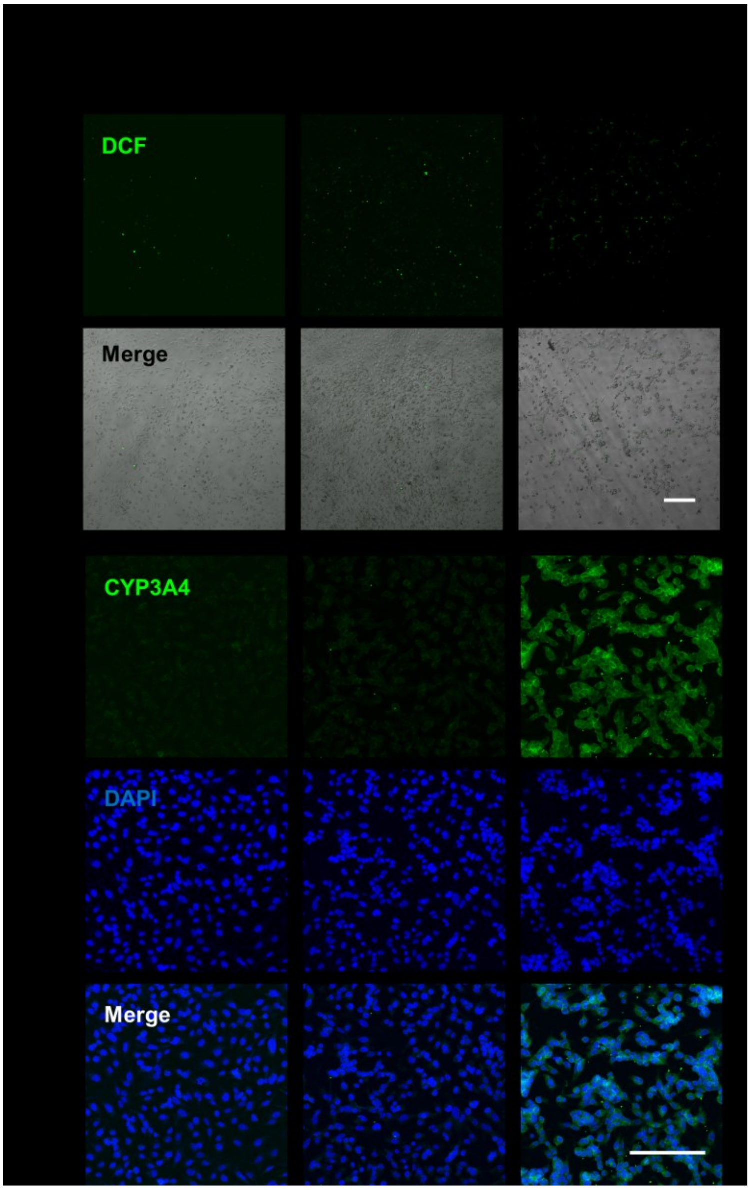


**Fig. S11.** (**A**) Representative fluorescence and bright-field merged images of cells in DOPC-DiO MP/NP T2, DOPC-DiO MP/NP T24 and DOPC-DiO MP/NP T72 groups, labeled by DCFH-DA. (**B**) Representative immunofluorescence images of CYP3A4 protein in DOPC-DiO MP/NP T2, DOPC-DiO MP/NP T24 and DOPC-DiO MP/NP T72 groups. Scale bar, 400 μm.

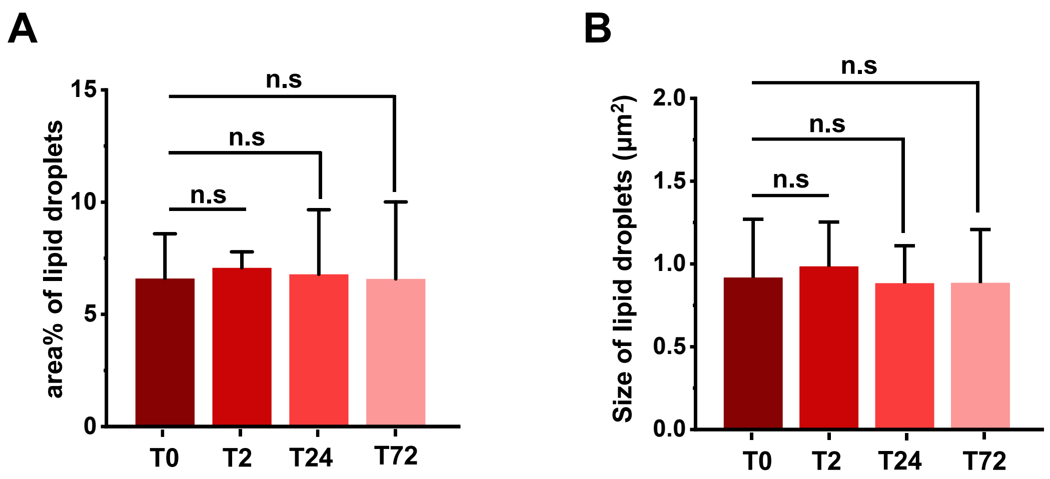


**Fig. S12.** (**A**) Size and (**B**) density of LDs in the MP/NP-treated cells of T0, T2, T24 and T72 groups. Values are mean ± SD (n=3). n.s, no significance.
